# High Performance protocol for ultra-short DNA sequencing using Oxford Nanopore Technology (ONT)

**DOI:** 10.1101/2025.01.10.632410

**Authors:** Lukas Žemaitis, Rūta Palepšienė, Simonas Juzėnas, Gediminas Alzbutas, Pierre-Yves Burgi, Thomas Heinis, Jérôme Charmet, Silvia Angeloni Suter, Martin Jost, Renaldas Raišutis, Friedrich Simmel, Ignas Galminas

**Affiliations:** Department of DNA data storage, Genomika, Kaunas, Lithuania; Ultrasound Research Institute, Kaunas University of Technology, Kaunas, Lithuania; Institute for Digestive Research, Academy of Medicine, Lithuanian University of Health Sciences, Kaunas, Lithuania; Department of Information and Communication Systems and Technologies, University of Geneva, Geneva, Switzerland; Department of Computing, Imperial College London, London, United Kingdom; School of Engineering HE-Arc Ingénierie, HES-SO University of Applied Sciences Western Switzerland, Neuchâtel, Switzerland; MABEAL GmbH, Graz, Austria; Department of Bioscience, TU Munich, School of Natural Sciences, Garching, Germany; Faculty of Natural Sciences, Vytautas Magnus University, Kaunas, Lithuania

**Author notes:** These authors contributed equally to this work (Lukas Žemaitis, Rūta Palepšienė).

## Abstract

In recent years, Oxford Nanopore Technologies (ONT) has gained substantial attention across various domains of nucleic acids’ research, owing to its unique advantages over other sequencing platforms. Originally developed for long-read sequencing, ONT technology has evolved, with recent advancements enhancing its applicability beyond long reads to include short, synthetic DNA-based applications. However, sequencing short DNA fragments with nanopore technology often results in lower data quality, likely due to a lack of protocols optimised for these fragment sizes. To address this challenge, we refined the standard ONT library preparation protocol to improve its performance for ultra-short DNA targets. Utilising the same core reagents required for conventional ONT workflows, we introduced targeted alterations to enhance compatibility with shorter fragment lengths. We then benchmarked these adjustments against libraries prepared using the standard ONT protocol. Here, we present a comprehensive, step-by-step protocol that is accessible to researchers of varied technical expertise, facilitating high-quality sequencing of ultra-short DNA fragments. This protocol represents a significant improvement in sequencing quality for short DNA fragments using ONT technology, broadening the range of possible applications.

**Graphical abstract:** 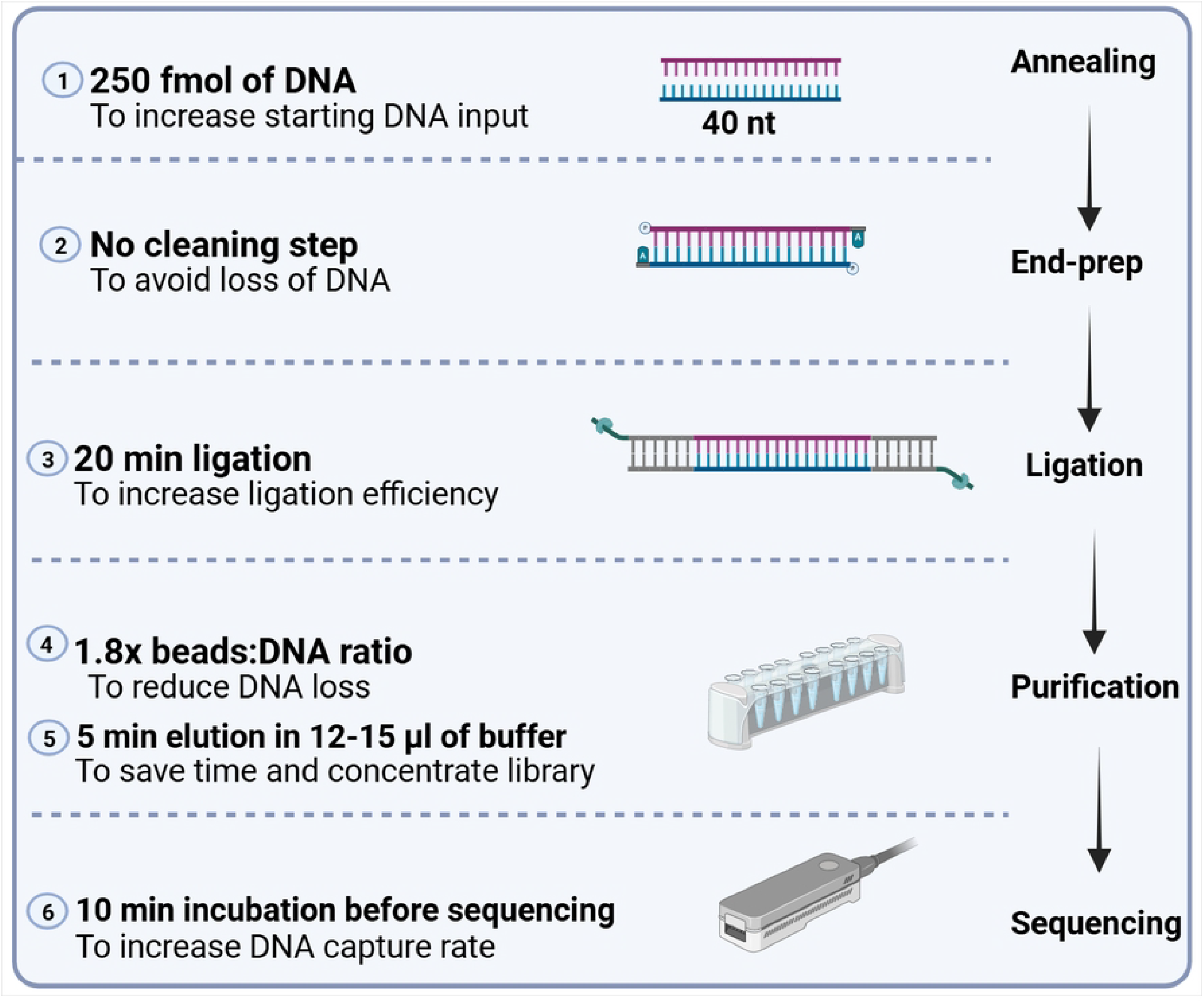

## Introduction

Oxford Nanopore Technologies has revolutionized sequencing by offering solutions that combine ease of operation, portability, accessibility, and rapid data acquisition. Unlike traditional sequencing technologies, ONT’s platform does not require elaborate sample preparation or complex sequencing infrastructure, allowing researchers to analyse DNA in real time and in various settings [1]. These appealing features have made nanopore sequencing increasingly popular across multiple domains. Beyond its widespread use in genomics and life sciences, there is growing interest in the application of nanopore sequencing in emerging fields such as DNA data storage [2], [3], DNA origami [4], and nanomaterials science [5], where DNA is utilized for both data encoding and as a structural component.

These emerging applications rely on short (< 100 bp) or ultra-short (20-40 bp) [6] synthetic DNA, a constraint imposed by the phosphoramidite chemistry typically employed in oligonucleotide synthesis [7]. The ONT long-read sequencing platform, however, encounters challenges when sequencing these oligonucleotides, often leading to substantial underutilisation of flow cells with reduced sequencing output and quality [8]. To counter these limitations, some strategies have focused on sample barcoding and multiplexing to maximise flow cell utilisation [9] while others attempted to concatenate ultra-short fragments into longer stretches, improving both efficiency and read quality [10]. Nevertheless, these strategies introduce additional steps, complicating and extending the library preparation process, and certain applications require native sequencing rather than concatenated forms. These challenges highlight the need for further optimisations to the current protocols to make ONT more effective for applications involving short oligonucleotide sequencing.

Here, we introduce an optimised, step-by-step protocol for sequencing DNA fragments as short as 40 base pairs (bp) using ONT. To develop this protocol, we systematically evaluated various modifications to the standard ONT protocols and identified six key steps that significantly enhance ONT performance for sequencing ultra-short DNA. We then benchmarked our protocol against the standard ONT protocol, demonstrating that by refining the standard Ligation Sequencing Kit V14, our approach achieves over ten times the sequencing output without the need for additional reagents apart from those used in standard library preparation.

The high reproducibility and streamlined nature of our protocol make it a compelling alternative to standard ONT protocol, particularly for applications involving short DNA fragments. Previously, the limited output for short fragments posed significant challenges, making the technology both inefficient and costly. Our approach, however, transforms this paradigm, offering a solution that overcomes these limitations. What is more, due to its simplicity and low complexity, this protocol can be seamlessly integrated into workflows across various laboratories, regardless of the users’ technical backgrounds. This broad applicability supports expanding ONT into research fields where the sequencing of ultra-short DNA fragments has traditionally posed challenges. By offering a significant boost in sequencing output and accuracy for short DNA fragments, this protocol has the potential to improve ONT usability and integrate it into new scientific domains, enhancing the quality of research outcomes.

## Materials and Methods

### DNA preparation

All experiments were performed using ultra-short DNA sequences of 40 bp. These sequences were synthesised as complementary ssDNA oligonucleotides using an on-site DNA synthesiser (Kilobaser) [11] and subsequently dissolved in 20 μL of Milli-Q water (Thermo Fisher Scientific). Concentration measurements were conducted using a Qubit 4 Fluorometer with the Qubit ssDNA assay kit (Thermo Fisher Scientific). Sequences used in the study were:

40_TOP 5’ – CTAACCCGACCTAATATGAACTCCTGCCTACAATGACCCT – 3’

40_BOT 5’ – AGGGTCATTGTAGGCAGGAGTTCATATTAGGTCGGGTTAG– 3’.

To obtain dsDNA fragments, 23 μM of 40_TOP and 40_BOT DNA oligonucleotides were combined with 1 μL of T4 ligase buffer (10x) (Thermo Fisher Scientific) and Milli-Q water, resulting in a final reaction volume of 10 μL. The reaction mixture was denatured at 95°C for 5 minutes, followed by gradual cooling at a rate of 5°C/min until reaching 25°C to promote annealing. After the reaction, the concentration of the dsDNA duplex was measured using the Qubit HS dsDNA assay kit (Thermo Fisher Scientific).

### Library preparation for ONT

Library preparation was performed using the Ligation Sequencing Kit with V14 chemistry (SQK-LSK114, ONT). Pre-annealed 40 bp fragments were subjected to library preparation by following the standard ONT protocol according to the manufacturer’s guidelines [12] (ACDE_9163_v114_revS_29Jun2022 protocol version) or a High-performance protocol, which is being proposed in this study. The detailed step-by-step library preparation method using the High Performance protocol is described and included for printing as S1 File with this article. In brief, an increased DNA input of 250 fmol of dsDNA duplex was phosphorylated at the 5’ end and adenylated at the 3’ end, following the manufacturer’s instructions. Without purification, ligation adapters were then attached to the processed samples using the Quick T4 DNA Ligation Module (New England Biolabs) for extended reaction time (20 mins). Following ligation, the samples were purified using AMPure XP beads with an increased beads-to-DNA ratio of 1.8x and subsequently eluted.

Samples prepared using the standard ONT protocol exhibited library concentrations below measurable limits; therefore, 10 μL of the prepared library was used for sequencing. In case of samples prepared with the High-Performance protocol, 8-10 μL (40±3 fmol) of the DNA library was loaded onto the flow cell to ensure adequate pore occupancy. This was followed by a 10-minute incubation period prior to initiating the sequencing experiment.

### Sequencing

Sequencing was conducted using R10.4.1 Flongle flow cells (ONT) and a MinION Mk1B sequencing device (ONT), adhering to the default parameters specified for the ligation sequencing kit. The sequencing runs were performed for a duration of 1.5 hours, with raw signal data collected in POD5 format and subjected to basecalling using Dorado (v0.7.2) basecaller with simplex v5.0 SUP (dna_r10.4.1_e8.2_400bps_sup@v5.0.0) basecalling model from ONT. The resulting sequences in FASTQ files were categorised into “passed” or “failed” based on their average quality scores (q-scores), which were calculated by employing fx2tab from SeqKit (v2.8.2) [13]. Only reads with q-scores above or equal to the threshold of 9, which is the default value for MinKNOW software, were included in subsequent mapping and clustering analyses.

### Bioinformatic Analysis

#### Mapping

Reads that passed the q-score threshold were aligned to a reference sequence (40_TOP) using the LAST (v1542) [14] aligner with local alignment mode and default parameters. To extract the number of mismatches (NM) and the CIGAR string, the alignments in MAF were converted to SAM format using maf-convert from the LAST toolkit. Sequences with 3 or fewer NM and an aligned length of 37 to 40 bases to the reference were considered mapped. To obtain error profiles from the mapped reads, a custom script was used to parse the CIGAR string to calculate substitution, deletion and insertion rates per position relative to the reference sequence, excluding hard-clipped bases.

#### Clustering

Raw sequencing reads with the adapters and an average quality score ≤9 or lengths shorter than 50 bp or longer than 300 bp were discarded. High-quality reads were dereplicated using VSEARCH (v2.22.1) [15]. The dereplicated reads were clustered using Swarm (v3.1.5) [16]with a local clustering distance threshold of d = 10. Clustered sequences were aligned using MAFFT (v7.525) [17]. Alignments were trimmed using TrimAl (v1.5.rev0) [18]. After trimming, sequences were reclustered using Swarm with the same parameters (d = 10), and representative sequences were selected. Representative sequences were compared against reference sequences using BLASTn (v2.15.0+) [19] with the blastn-short task. Data analysis and visualisation were performed using custom R scripts in R (v4.3.3). The analysis pipeline was managed using Snakemake (v8.11.3) [20].

### Statistical analysis

Statistical analysis was performed using R (v4.4.0) [21] and ggplot2 (v3.5.1) [22] was used for visualization. Data are presented as the mean ± standard error (standard deviation divided by the square root of the sample size) or as the median, including the first and third quartiles and a 1.5× interquartile range. Statistical significance between the two sample groups was evaluated using an unpaired Student’s t-test with a “greater” one-sided alternative hypothesis.

## Expected results

Recent advancements in Oxford Nanopore Technologies suggest that it is possible to sequence DNA fragments as short as 20 bp. Yet, previous studies have highlighted a notable decline in ONT performance with short DNA fragments [8], [10] which may be further aggravated when targeting ultra-short DNA. Therefore, we selected an ultra-short DNA fragment of 40 bp and aimed to address key questions: i) evaluating the performance of ONT for ultra-short DNA fragments; ii) determining how adjustments of the current protocols might enhance sequencing quality for such fragments.

Our preliminary pilot experiments (data not shown) confirmed that standard ligation-based library preparation protocol provided by the ONT produce a low read count and diminished quality scores. Recognising the potential of this technology, we refined the standard protocol to make it more suitable for ultra-short fragment sequencing. This article describes an optimised approach, termed the High-Performance Protocol, which was rigorously evaluated through benchmarking against the standard ONT library preparation protocol, focusing on key sequencing performance metrics, including sequencing throughput, quality, mapping and error rates. The study design, alongside the methodological distinctions between the protocols, is outlined in Fig. 1A.

**Fig 1.**
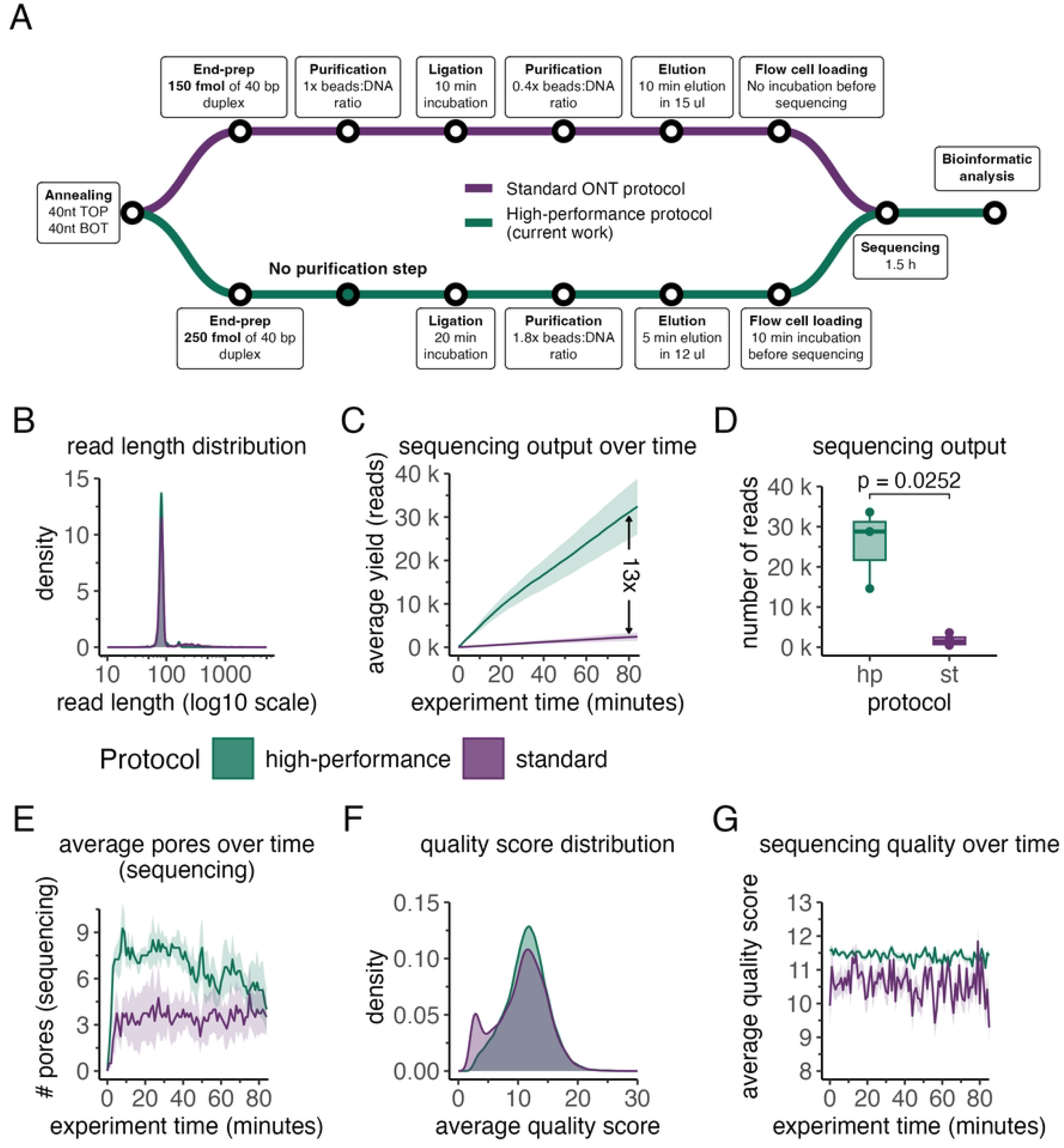
The high-performance enhances sequencing efficiency compared to standard library preparation in ONT nanopore sequencing. **A)** Detailed schematics of the experimental design and the key differences in library preparation methods to illustrate the workflow and methodology. **B)** Sequencing read length distributions for libraries using standard (purple) and high-performance (green) protocols. **C)** Average cumulative throughput (reads) and standard error (SE; y-axis) versus experiment duration (sequencing) (x-axis). D) Number of “passed” reads (q-score ≥ 9; y-axis) generated using high-performance and standard library preparation protocols (x-axis). The boxplots show the median (centre line), first and third quartiles (lower and upper hinges), and 1.5× interquartile range (lower and upper whiskers). Statistical significance between the protocols was evaluated using an unpaired Student’s t-test with a one-sided “greater” alternative hypothesis. **E)** Number of pores in the sequencing state (y-axis) throughout the experiment duration (x-axis). The line graph shows the mean ± SE of pore numbers in the sequencing state for both high-performance and standard libraries. **F)** Q-score distributions of libraries prepared by standard (purple) and high-performance (green) protocols. **G)** Average q-scores with SE of sequencing reads (y-axis) over the experiment time (x-axis) generated by both the High-performance and standard libraries. The results represent three (N = 3) and four (N = 4) independent sequencing experiments by using the High-performance and standard ONT protocol, respectively.

Our primary goal was to determine whether overall sequencing quality (generated output and read quality) varies between protocols. Analysis of fragment length distributions revealed a peak at approximately 85 bp for both the standard and High-Performance protocols, aligning with the intended length of the annealed DNA duplex in combination with attached ONT sequencing adapters (Fig. 1B). This consistency shows that both protocols successfully prepared libraries of the intended fragment size. A notable distinction between protocols emerged in output efficiency: the High-Performance protocol yielded approximately 13 times more reads than the standard protocol (Fig. 1C). Specifically, the High-Performance protocol produced an average of 33252 raw reads, compared with 2551 raw reads for the standard protocol over the same sequencing period, underscoring its superior throughput.

Reads were then classified based on quality scores into high-quality and low-quality categories as described in Methods and Materials section. It was observed that both protocols yielded predominantly high-quality reads, with approximately 77% (25669 reads) and 68% (1740 reads) of total reads classified as high-quality for the High-Performance and standard protocols, respectively (Fig. 1D). The statistically significant difference (p = 0.0286) in the number of high-quality reads between the protocols primarily reflects the substantially higher initial read count generated by the High-Performance protocol. However, these results also indicate that the High-Performance protocol can achieve a marked increase in read counts without compromising read quality.

To investigate the underlying cause of the substantially higher read count observed with the High-Performance protocol, we hypothesised that it may be attributed to the amount of DNA input loaded onto the flow cell. As outlined in the Materials and Methods section, the precise quantification of DNA was not feasible for the standard ONT protocol due to the low concentration. In contrast, the High-Performance protocol consistently produced purified DNA libraries with measurable concentrations, of which ∼40 fmol, were loaded for sequencing. The higher concentration of DNA molecules would be expected to increase pore occupancy, thus generating a higher read count [23]. To test this hypothesis, we monitored the number of actively sequencing pores throughout the experiment (Fig. 1E). Consistent with our hypothesis, the High-Performance protocol maintained, on average, more than twice the number of pores in the active state as compared to the standard protocol. In addition to the higher DNA concentration, the High-Performance protocol includes an extra 10-minute incubation period before sequencing. This step may provide more time for tether molecules to bring DNA closer to the membrane, improving the DNA capture rate by the nanopore [24].

As the quality score of reads is as critical a metric as overall output, we conducted a more detailed analysis. A comparison of the average read quality scores between protocols revealed that both protocols produced reads with similar quality scores, centred around 11 (q-score=11) (Fig. 1F). However, the distribution patterns of read quality differed. The High-Performance protocol generated a relatively low proportion of reads with extreme quality values (q ≤ 5 or q ≥ 15), with most reads displaying intermediate quality, as expected for such short fragments. In contrast, the standard ONT protocol demonstrated a marked increase in reads with low quality scores of q ≤ 5. This discrepancy may be partially attributed to the generation of reads with highly variable quality scores over the course of the experiment (Fig. 1G) and to sequencing artifacts, which are unexpectedly long reads exhibiting low sequence diversity (Supplementary Figure 1). This was mostly observed in samples prepared using standard ONT protocol while High-Performance protocol maintained relatively stable quality scores and read length throughout the sequencing period. The factors contributing to this variability in samples prepared according to the standard protocol are not yet fully understood and will require further study.

Previous studies have indicated that ONT typically generates error rates of approximately 3-10% in case of long reads (>1 kb) [25], [26]. This error rate is expected to increase with ultra-short DNA fragments, potentially due to the rapid translocation speed of DNA through the nanopore which can hinder the precise interpretation of signals and accurate base identification during basecalling [1]. Consequently, in the next phase of this study, we evaluated the sequencing accuracy and compared potential differences between samples prepared using the standard and High-Performance protocols.

To conduct this analysis, we aligned high-quality (q-score ≥ 9) reads to a reference sequences and assessed accuracy from multiple perspectives. As a first step, we examined the average length of the mapped reads (Fig. 2A). Notably, the High-Performance protocol produced a substantially higher number of full-length reads compared to the standard protocol, which generated relatively fewer full-length reads. This difference suggests that the High-Performance protocol may provide better coverage of the target sequence, improving downstream analyses.

**Fig 2.**
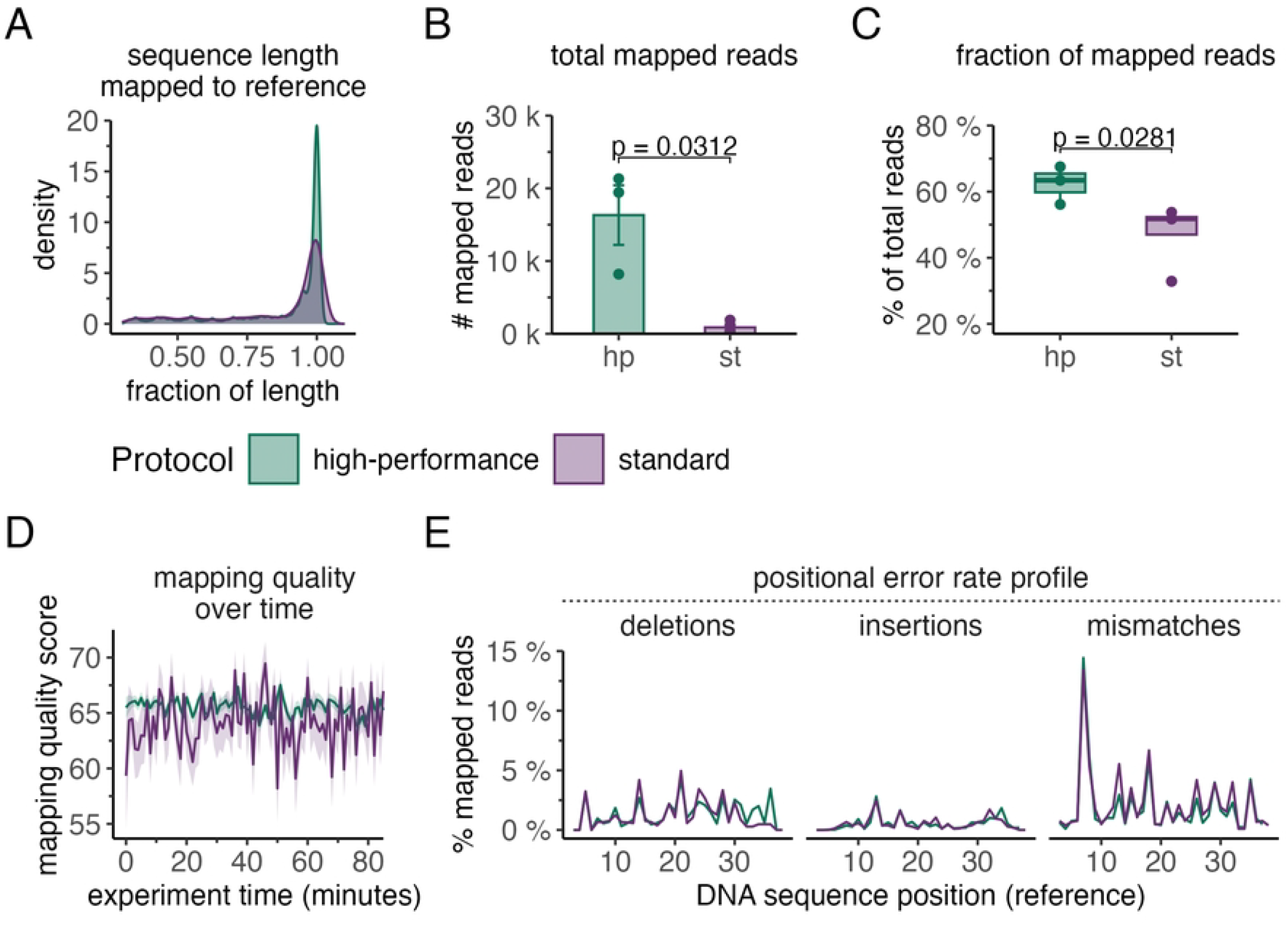
Sequencing reads generated by the high-performance protocol yield a higher proportion of correct, full-length sequences. **A)** Probability densities (y-axis) of mapped sequence lengths relative to the reference length (x-axis) for libraries prepared with both the standard protocol (purple) and the high-performance protocol (green). **B)** Number of reads (y-axis) mapped to the reference sequence with 3 or fewer mismatches for data generated by high-performance and standard library preparation protocols (x-axis). The bar plot displays the mean ± SE of reads numbers. **C)** Fraction of mapped reads relative to the total number of passed reads (y-axis) for sequencing libraries prepared using high-performance and standard library preparation protocols (x-axis). The boxplots display the median (centre line), first and third quartiles (lower and upper hinges), and 1.5× interquartile range (lower and upper whiskers). **D)** Average mapping-quality (MAPQ) values ± SE of passed sequencing reads (y-axis) over the experiment duration (x-axis) generated by both the high-performance and standard libraries. **E)** Positional fractions of deletions, insertions and mismatches (y-axis) with respect to the reference sequence (x-axis) for libraries prepared by high-performance (green) and standard (purple) library preparation protocols. The line graph displays the mean values for the fraction of total passed reads for the high-performance and standard libraries. The results represent three (N = 3) and four (N = 4) independent sequencing experiments by using the High-performance and standard ONT protocol, respectively.

Further distinctions between the protocols became evident when quantifying the total number of mapped reads (Fig. 2B). The High-Performance protocol yielded significantly more mapped reads than the standard protocol (p = 0.003), averaging 16320 reads compared to 874. This difference largely reflects the higher total number of high-quality reads generated by the High-Performance protocol. However, when we evaluated the mapping fraction (i.e., the proportion of mapped reads relative to the total number of high-quality reads), additional factors emerged. The samples prepared by the High-Performance protocol achieved a notably higher mapping fraction of approximately 60%, compared to only about 45% for the standard protocol, indicating significantly improved mapping efficiency. To test If this discrepancy might be related to overall lower quality scores in reads generated from samples prepared with the standard ONT protocol (as indicated in Fig. 1F and G) we analysed mapping quality over all 85-minute sequencing periods for each protocol (Fig. 2D). In accordance with our observation, mapping quality scores differed significantly between protocols: the High-Performance protocol maintained consistently high mapping quality, while the standard protocol exhibited fluctuations of greater amplitude. Positional error rate profiles were also compared between protocols (Fig. 2E), revealing similar overall error rates, with only a slightly higher error rate observed in reads generated by the standard protocol.

Finally, we investigated the structural composition of the reads using sequence clustering analysis. As presented in Fig. 3, reads from both protocols exhibited the expected structure, with the target sequence centred and flanked by adapter fragments. In the case of samples prepared using the High-Performance protocol (Fig. 3 A), the largest cluster comprised 43% of the total reads (32936 reads), while the second-largest cluster accounted for 27% (21130 reads). The largest cluster contained the correct insert in the forward orientation (plus strand), whereas the second-largest cluster corresponded to the same insert just in reverse complement orientation (minus strand). The data obtained from the standard protocol (Fig. 3 B) revealed a similar clustering pattern, but with reduced structural homogeneity: the largest cluster, presented the 35% of the total reads (2361 reads), corresponding to the plus strand, while the second-largest cluster accounted for 20% (1309 reads) and corresponded to the minus strand. These results highlight the structural homogeneity differences between the two protocols.

**Fig 3.**
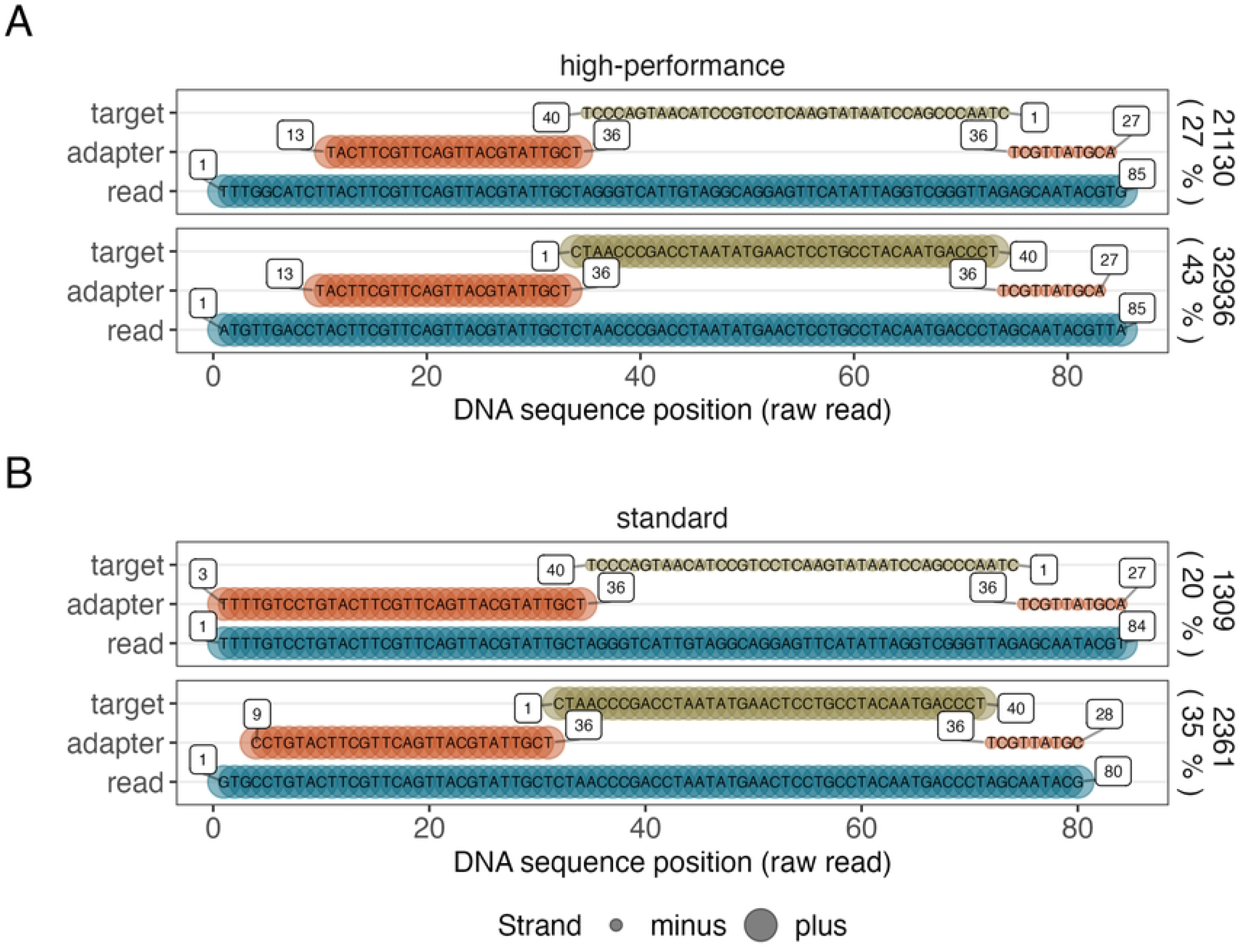
Clustering analysis reveals that high-performance protocol yields a higher fraction of reads containing the correct sequence. The graph represents the top two most abundant consensus sequences in libraries prepared by high-performance **(A)** and standard **(B)** library preparation protocols. Reads were aligned with the target sequence (green) flanked by adapter fragments (dark and light orange). Numbers within the bars indicate specific sequence positions. The proportions of reads in the largest and second-largest clusters are displayed to the right of each panel, alongside the total read counts for each cluster. The x-axis represents the DNA sequence position of raw reads, while strand orientation (plus or minus) is indicated by line thickness.

This clustering distribution suggests that the High-Performance protocol was more effective in preserving the structural integrity of the target sequence across its full length, as evidenced by the higher proportion of reads assigned to the dominant cluster. The increased homogeneity observed with the High-Performance protocol likely results from its ability to generate a higher proportion of full-length reads (Fig. 2A), which contributes to the observed clustering patterns.

## Conclusions

This study presents a refined High-Performance protocol, specifically designed to optimise sequencing of ultra-short DNA fragments. Relying on the same conventional reagents as those used in standard ONT protocols, the High-Performance protocol delivers over a tenfold increase in sequencing yield, while maintaining stable sequencing quality and improving mapping efficiency for fragments as short as 40 bp.

The protocol is also applicable to longer DNA fragments (> 40 bp), although with possibly reduced efficiency due to its underlying principles such as the limited effect of increased bead concentration on DNA recovery as fragment length increases. Further research is needed to evaluate its performance across different fragment lengths comprehensively.

Overall, this study highlights the advantages of the High-Performance protocol, demonstrating its potential for a wide range of applications previously considered incompatible with nanopore sequencing technology. This streamlined approach expands the capabilities of ONT sequencing, opening new possibilities for its use in fields focused on short and ultra-short DNA analysis.

## Authors’ contributions

**Conceptualization**: Rūta Palepšienė, Simonas Juzėnas, Gediminas Alzbutas, Ignas Galminas, Lukas Žemaitis

**Data Curation:** Simonas Juzėnas, Gediminas Alzbutas

**Formal Analysis:** Simonas Juzėnas, Gediminas Alzbutas

**Funding Acquisition:** Ignas Galminas, Lukas Žemaitis, Pierre-Yves Burgi, Thomas Heinis, Jérôme Charmet, Silvia Angeloni Suter, Martin Jost, Renaldas Raišutis, Friedrich Simmel

**Investigation:** Rūta Palepšienė, Lukas Žemaitis

**Methodology:** Rūta Palepšienė, Lukas Žemaitis

**Project Administration:** Ignas Galminas

**Resources:** Ignas Galminas, Lukas Žemaitis

**Supervision:** Ignas Galminas, Lukas Žemaitis

**Software:** Simonas Juzėnas, Gediminas Alzbutas

**Validation:** Rūta Palepšienė

**Visualization:** Simonas Juzėnas

**Writing – Original Draft Preparation:** Rūta Palepšienė, Simonas Juzėnas, Gediminas Alzbutas

**Writing – Review & Editing:** Rūta Palepšienė, Simonas Juzėnas, Gediminas Alzbutas, Ignas Galminas, Lukas Žemaitis, Pierre-Yves Burgi, Thomas Heinis, Jérôme Charmet

## Data availability

The data and code associated with this study are available on the GitHub repository (https://github.com/genomika-lt/high-performance) and archived on Zenodo (https://zenodo.org/records/14244834).

## Supporting information

**S1 File. Step-by-step High-performance protocol**

**S1 Fig. The high-performance protocol produces fewer sequencing artifacts than the standard library preparation method. A)** Average read length of sequences produced over time are more homogenous for the libraries prepared by high-performance protocol. The line graph shows the average read length with standard error (SE) for libraries prepared using the standard and high-performance protocols. **B)** As shown in the boxplots, libraries prepared using the standard (st) library preparation protocol have a higher fraction of reads with a length of 1 kb or more compared to those prepared using the high-performance (hp) protocol. Of note, sequencing artifacts in this case refer to unexpectedly long reads exhibiting low sequence diversity. The results represent three (N = 3) and four (N = 4) independent sequencing experiments by using the High-performance and standard ONT protocol, respectively.

